# Age-dependent selection on MHC class 2 variation in a free-living ruminant

**DOI:** 10.1101/2020.03.25.008565

**Authors:** Wei Huang, Kara L Dicks, Jarrod D Hadfield, Susan E Johnston, Keith T Ballingall, Josephine M Pemberton

## Abstract

Genes within the major histocompatibility complex (MHC) are the most variable identified in vertebrates. Pathogen-mediated selection (PMS) is believed to be the main force maintaining diversity at MHC class I and II genes, but it has proven hard to demonstrate the exact PMS regime that is acting in natural populations. Demonstrating contemporary selection on MHC alleles is not trivial, and previous work has been constrained by limited genetic tools, low sample sizes and short time scales and has sometimes involved anticonservative statistical approaches. Here, we use appropriate statistical approaches to examine associations between MHC variation and several fitness measurements including total fitness (lifetime breeding success) and five fitness components, in 3400 wild Soay sheep (Ovis aries) monitored over their lifetimes between 1989 and 2012. We found haplotypes C and D were associated with decreased and increased male total fitness respectively. In terms of fitness components, juvenile survival was positively associated with haplotype divergence. Of the eight MHC haplotypes (A-H), haplotypes C and F were associated with decreased adult male breeding success and decreased adult female life span respectively. Consistent with the increased male total fitness, haplotype D, which is the rarest, has increased in frequency throughout the study period. Our results suggest that contemporary balancing selection is acting on MHC class II genes in Soay sheep and that different selection mechanisms are acting between juveniles and adults.

## Introduction

The vertebrate immune system has evolved to protect the host from infection through a variety of innate and adaptive immune mechanisms. Many of the proteins that control the specificity of immune function are the products of genes that are amongst the most diverse within vertebrate populations. Such diversity is believed to have evolved in response to pathogen diversity. Investigating the mechanisms shaping the diversity of immune genes has been a major focus in evolutionary biology (1). Genes located within the major histocompatibility complex (MHC) encode proteins that play important roles in the vertebrate immune system and have been intensively studied across many species, including model organisms (2–4). MHC molecules, the products of MHC class I and class II genes, present peptide fragments to antigen-specific receptors on T cells to induce adaptive immune responses. MHC class I molecules present peptides derived from intracellular pathogens such as viruses to CD8 +ve T cells, while MHC class II molecules present peptides derived from extracellular pathogens such as bacteria and parasites to CD4 +ve T cells (5).

MHC class I and II genes are the most polymorphic loci identified in vertebrate populations with hundreds of alleles described in many species (6, 7). Pathogen-mediated balancing selection is thought to be the principal driver maintaining such high levels of allelic diversity (8–11). However, the specific selection mechanisms which act to maintain high levels of MHC allelic diversity are not clear (3, 12, 13). Three models have been proposed to explain how PMS acts to maintain such diversity:

1. *Heterozygote advantage*: individuals which are more heterozygous across the MHC can respond to a greater variety of pathogens. As a result, variation at MHC loci is selected and maintained (14). In addition, divergent allele advantage has been proposed as an extension to heterozygote advantage. Here, individuals with a greater functional divergence between alleles should have a selective advantage because their MHC molecules can bind a broader range of antigens (15).
2. *Rare-allele advantage*: there is strong selection on pathogens to evade immune protection provided by the most common host MHC alleles. Therefore, rare alleles have a selective advantage. However, the advantage disappears as the frequency of a protective MHC allele increases and is absent at equilibrium. The arms race between hosts and pathogens sets up a cyclical process causing MHC alleles and pathogens to fluctuate, meaning that MHC diversity is maintained by negative frequency-dependent selection (16–18). It should be noted that since rare alleles appear most frequently as heterozygotes, this can manifest as heterozygote advantage for specific alleles under some circumstances.
3. *Fluctuating selection*: a changing environment results in variation in the abundance of different pathogens in space and time, which generates directional selection in different host subpopulations and/or at different time points. Thus, different alleles will be favoured in different subpopulations or at different times, meaning MHC diversity is maintained as a consequence (19, 20). Unlike rare-allele advantage the change in selection coefficients in time and/or space is not driven by coevolution in the pathogens.

To test each selection model we can examine associations between MHC diversity and parasite presence or parasite load; different patterns of associations can be explained as a result of different selection mechanisms (13). A number of studies on MHC genes in natural populations have examined associations between MHC variation and parasites, often with conflicting results. Some studies found that MHC heterozygosity or certain MHC alleles confer resistance to parasite infection (21–23), while other studies found no association between MHC alleles and parasite load (24). However, such studies may be limited by incomplete data on the diversity and abundance of the parasite populations present and parasites that strongly select for MHC diversity may have been missed. As a consequence, quantifying relationships between MHC genes and pathogens may not have been able to explicitly elucidate which selection mechanisms were operating, although some experimental studies have managed to demonstrate a clear mechanism by examining associations between MHC and pathogens (25, 26). An alternative means to test selection on MHC genes would be to examine MHC-dependent fitness patterns (4, 27).

As with studies examining associations between MHC and parasites, MHC-fitness associations can also indicate different selection mechanisms. Under heterozygote advantage, we predict a positive association between MHC heterozygosity and fitness. Under rare-allele advantage, if MHC alleles are not in equilibrium, we may detect selection favouring specific rare alleles and probably also disfavouring specific common alleles. Under fluctuating selection, we would also observe specific alleles being favoured or disfavoured by selection. But unlike rare-allele advantage, this should occur regardless of their frequency. However, it is hard to distinguish rare-allele advantage and fluctuating selection by examining MHC-fitness associations (13). Because measuring fitness is generally challenging except in some long-term individual-based studies of natural populations, only a few studies have tested associations between fitness and MHC variation in the wild, again with mixed results. First, some MHC studies in wild populations have found a positive association between MHC heterozygosity and fitness. For instance, the number of MHC alleles was found to be positively associated with increased apparent survival (return rate from initial year of capture) in common yellowthroats (Geothlypis trichas) (28). Similarly, Banks et al. found an association between MHC heterozygosity and survival in a natural population of mountain brushtail possums (Trichosurus cunninghami). Second, some studies have reported associations between fitness and specific MHC alleles or supertypes (clusters of MHC alleles with similar physicochemical properties at their antigen-binding sites). For example, Lukasch et al. (30) found that a specific MHC allele was associated with higher survival of house sparrow fledglings (Passer domesticus). Another study of great tits (Parus major) using mark-recapture data identified three MHC supertypes associated with fitness components, one with increased survival, one with increased lifetime reproductive success and another one with decreased lifetime reproductive success (31). Kloch et al. (32) also found that two MHC alleles from different phylogenetic clusters were associated with winter survival in root voles (Microtus oeconomus). Finally, there are some studies where no associations are detected. In collared flycatchers (Ficedula albicollis), no association between MHC variation and lifetime reproductive success was found (24). Similarly in house sparrows (Passer domesticus), no significant associations between MHC variation and survival were identified (33).

A complication in studies of fitness association is that associations may differ between individuals of different sex, age or nutritional status, presumably due to variation in their ability to counter the threat from pathogens. For example, in a population of black-legged kittiwake (Rissa tridactyla), the association between MHC diversity and fitness was only found in second-hatched female chicks suggesting sex and hatching order could modulate MHC-fitness association (34). In a previous study of Seychelles warbler (Acrocephalus sechellensis), a positive association with MHC diversity was only found in juvenile survival but not in adult survival (35). However, the impact of age on MHC-fitness associations has been rarely studied because of the limitation of study length and scale in wild populations. Thus, longitudinal studies with fitness measurement through all life stages of study individuals are required to study MHC-dependent fitness effects.

Apart from the difficulty in measuring fitness to study selection on MHC genes, many previous studies of selection on MHC genes are limited by the depth and quality of the genetic data available. Within a typical mammalian MHC region, multiple closely linked and duplicated genes are present which are inherited together as a haplotype. With the development of next generation sequencing (NGS) technologies, we are now able to cost-effectively genotype large numbers of samples (36, 37). However, in species where the MHC region is poorly characterised such as in fish or passerine birds, it is difficult to assign a sequence to an individual locus or identify if a locus is functional. Without locus-specific genotyping, multiple loci may co-amplify, which will impact subsequent testing for associations between MHC diversity and fitness. For example, heterozygote advantage cannot be demonstrated without knowing that alleles are alternatives at the same locus. Even if locus-specific genotyping is possible, many studies have focused on a single MHC locus and ignored linkage with other MHC loci (29, 32, 38). Locus-specific genotyping and characterisation of haplotypes are desirable when testing for associations between fitness and MHC diversity in wild populations.

Lastly, statistical analyses of selection on MHC variation could be more precise for controlling confounders than is often deployed. Measures of parasite load and fitness are quantitative traits likely to be determined by both environmental factors and multiple genetic factors, including MHC variation. As such, it is possible that studies of contemporary selection may confound MHC similarity with genetic relatedness, so an MHC effect could be due to variation elsewhere in the genome rather than the MHC itself. Selection on MHC genes in natural populations has mostly been assessed in long-term individual-based studies which include a lot of genetically related individuals (31, 35). To our knowledge, few MHC studies in natural populations have been conducted in a quantitative genetic framework. In addition, given the ubiquity of inbreeding depression, any analysis of selection should include an estimate of individual inbreeding to ensure MHC effects are not confounded with genome-wide inbreeding effects. However, only a few previous studies, e.g. (39, 40) have accounted for inbreeding.

Here, we present the first investigation of contemporary selection acting on the MHC in a free-living population that overcomes the issues raised above. The Soay sheep (Ovis aries) population on the island of Hirta, St Kilda is one of the most intensively studied wild animal populations in the world (41). Since 1985, individual Soay sheep living in the Village Bay study area have been followed from birth, through all breeding attempts, until death. In addition, using a combination of observation and SNP genotypes, a multigenerational pedigree for the population has been assembled which includes annual and lifetime measures of fitness for both sexes (42). Previous work characterised diversity at the MHC class II DRB1 gene using an internal microsatellite and identified associations between MHC genotype, fitness and parasite infection (43, 44). However, the only fitness measure studied was juvenile survival. Another study using the same genetic dataset of MHC-linked microsatellite alleles found higher temporal and spatial variation in these loci than in putatively neutral microsatellite loci in other parts of the genome, suggesting fluctuating selection on MHC genes (45). Recently, using sequencing-based genotyping of functional MHC class II DR and DQ loci we have shown that eight MHC haplotypes are segregating in the study population (46). Using a panel of 13 SNPs, MHC class II diplotypes (i.e. haplotype complement) were characterised for 5349 individuals sampled between 1989 and 2012 (47). Diplotypes were in Hardy-Weinberg equilibrium both within cohorts (cite Kara BioRxiv MS) and within the population alive each year (47). Here, we use this data to investigate associations between MHC variation and fitness. We test for associations between (1) MHC variation and total fitness, defined as lifetime breeding success of each individual and (2) MHC variation and five fitness components. We fit MHC variation including divergence, heterozygosity and individual haplotypes within quantitative genetic models in order to distinguish the effect of MHC haplotypes from genome-wide effects. Finally (3), we perform gene-drop simulations to examine whether haplotype frequencies have changed more than expected from a neutral process.

## Methods

### Study system

Soay sheep have lived virtually unmanaged on the island of Soay, in the St. Kilda archipelago, for thousands of years. In 1932, 107 Soay sheep were introduced to the larger neighbouring island of Hirta and have been living there unmanaged since. A previous study demonstrated that an introgression event between Soay sheep and a more modern breed occurred approximately 150 years ago (48). From 1985, a long-term study has been conducted on the sheep resident in the Village Bay area of Hirta to investigate ecological and evolutionary questions (41). The vast majority of lambs are born in April or May of each year. At their first live capture, lambs are ear-tagged to enable life-long identification and discs of ear tissue removed in the tagging process are used for DNA extraction. Parentage is inferred for each individual using a subset of 315 unlinked SNPs derived from the Illumina Ovine 50K SNP array, on which most individuals alive since 1990 have been genotyped (49). In addition, the R package Sequoia (50) was used to cluster half-siblings which share a parent that has not been genotyped. In cases where no SNP genotypes were available, a small number of parentage inferences were made using field observations (for mothers) or a previous microsatellite genotype approach (51). The pedigree used here includes 7447 individuals, with 7014 paternal links and 6229 maternal links. This study investigated associations between MHC variation and fitness measurements defined as follows:

- Total fitness, the number of offspring an individual produced over its lifetime;
- Juvenile survival (JS), a binary variable indicating whether a lamb survived from birth until May 1st of the following year.
- Adult annual survival (adult AS), a binary variable indicating whether an adult survived or not in a given calendar year; an adult is an animal that survived to age 1.
- Adult life span (adult LS), the age of an adult when it died;
- Adult annual breeding success (adult ABS), the number of offspring an adult produced in a given calendar year;
- Adult lifetime breeding success (adult LBS), the number of offspring an adult produced over its lifetime;

### MHC Class IIa Haplotyping

The genetic data used in this study were obtained from a previous study (46, 47). In that study the seven expressed loci (DRB1, DQA1, DQA2, DQA2-like, DQB1, DQB2 and DQB2-like) within the MHC class IIa region were characterised in 118 selected Soay sheep using sequencing-based genotyping. A total of eight MHC haplotypes were identified and named A to H. These haplotypes were confirmed in an additional 94 individuals that were unrelated based on the pedigree. A panel of 13 SNPs located in the region of MHC class IIa haplotypes was then selected to impute the eight MHC class IIa haplotypes in 5951 Soay sheep genotyped using Kompetitive Allele-specific PCR (KASP). This panel included 11 SNPs known to be variable in Soays based on data from the Illumina Ovine infinium HD chip and two other SNPs identified within the DQA1 gene (52). After quality control, the diplotypes of 5349 individuals that lived in the study area between 1985 and 2012 were identified. As cohort effects were fitted in the statistical analyses, we excluded animals born before 1989 since too few were genotyped to be representative of their birth cohort. We also removed the cohort born in 2010 which had a high genotyping failure rate in the KASP genotyping assay, all animals which died as foetuses and any individuals subjected to experimental treatments which may have affected their survival or breeding performance (e.g. anthelmintic boluses). After these exclusions, a total of 3440 individuals (1750 males and 1650 females) remained for statistical modelling of total fitness and JS (Table 1). The frequency of the rarest haplotype (D) was 3.02%. With the exception of animals homozygous for haplotype D (N=5), moderate numbers of individuals (N>20) of each diplotype are represented (Supplementary 1). For each individual successfully diplotyped, MHC divergence was measured by the proportion of the amino acid sequence that differed between the two MHC haplotypes (p-distance)(53) (Supplementary 2).

**Table 1.**
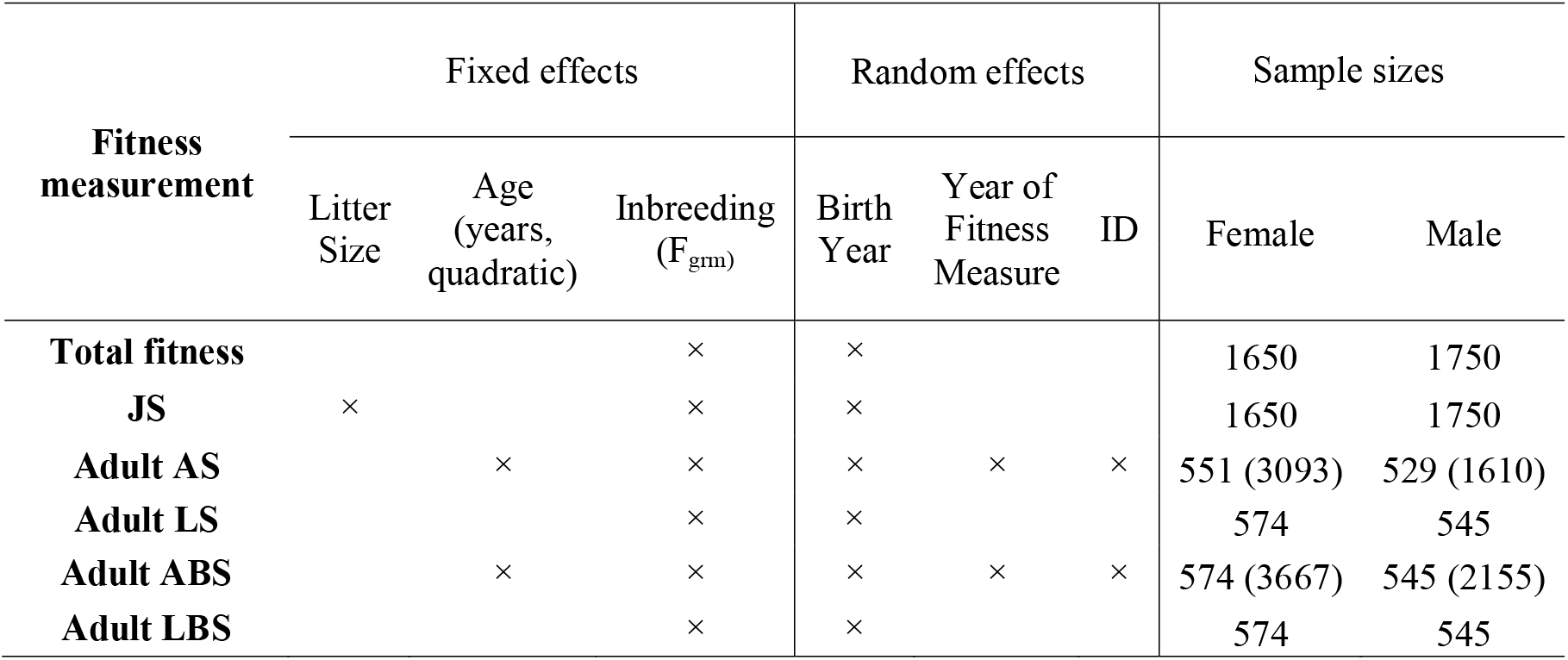
Fixed and random effects fitted in null model and sample sizes, for each fitness measurement. Sample sizes are shown by number of individuals (records).

### Statistical analysis

We used generalized linear mixed models (GLMMs) to study the associations between MHC haplotypes and fitness measurements. The primary models were animal models (AMs) that include a covariance structure proportional to the pedigree relatedness matrix, but we also repeated all tests without pedigree information. In null models, we fitted a set of fixed and random effects for each fitness measurement (Table 1) including genomic inbreeding (Fgrm) as a fixed effect, as previous studies demonstrated inbreeding depression in this population (54, 55). None of the fitness measurements in our study had a normal distribution, so we used Markov chain Monte Carlo techniques implemented in MCMCglmm v 2.28 (56) to run models.

To examine whether there were associations between MHC variation and fitness measurements, we fitted MHC heterozygosity, MHC divergence and individual MHC haplotypes simultaneously into the null model. Each haplotype was fitted as dosage (0, 1 or 2 copies) as a fixed effect in all the models. We performed a Wald test using the posterior mean and the posterior covariance matrix to test whether MHC haplotypes explained variation in fitness. In these models, Haplotype H was treated as a reference so the results for individual haplotypes from our models are relative to haplotype H. When a Wald test for haplotype differences was significant, we conducted an additional analysis (Supplementary comparing the estimate for each haplotype with the mean of the estimates for all other haplotypes, to identify individual haplotype effects.

Finally, we used models with different error distributions to combat the heterogeneity of ontogenies and distributions of fitness measurements (Supplementary 3). We used probit regression to run a unisex model for JS. As the ontogenies and distributions of total fitness and adult fitness components were different between the two sexes, we used separate models for males and females. For both males and females, we used zero-inflated Poisson models for total fitness, probit regression for adult AS and Poisson regression for adult LS and LBS. As females can only have up to three offspring each year while males had between 0 and 24, we used multiple category probit (ordinal) regression for female ABS and Poisson regression for male ABS. All the models were run for 200,000 iterations in R v3.5.2 (57).

### Gene-drop analysis

We performed gene-drop simulations with a custom script in R v3.5.2 (57) to model the expected frequency change of MHC haplotypes under genetic drift given the pedigree structure. Hereafter, the first four birth cohorts (1989 to 1992) are defined as the “founder” cohorts, whereas all subsequent cohorts are defined as “simulated” cohorts (1993 to 2012). A total of 5000 simulations were conducted as follows. Individuals in the founder cohorts were assigned their observed diplotypes; founder cohort individuals with unknown diplotypes were assigned one by either: (a) sampling a haplotype from each parent assuming Mendelian segregation; or (b) sampling a haplotype with the probability of the observed haplotype frequencies in the same birth cohort when one or more parents were unknown. In the simulated cohorts, each individual diplotype was sampled using approaches (a) and (b) above. By this method, simulated diplotypes were generated for each individual in the pedigree born after 1992. Using these data with the record of birth year and death year of each individual, we compared the observed and simulated changes in MHC haplotype frequency for the standing population (all the individuals living in a single year), using a linear regression with year (1993 to 2012) as the predictor variable. In cases where the observed slope fell within the top or bottom 2.5% of simulated slopes, the allele frequency was deemed to have been subject to directional selection.

## Results

### Associations between MHC and fitness measurement

The results of models with and without pedigree information were largely similar (Supplementary 5 to 7). Here we report the results of animal models, i.e. including the pedigree as a random effect.

#### Total fitness

There was no association between MHC heterozygosity or MHC divergence and total fitness in either sex (Figure 1A). In the Wald test for haplotype effects a significant association was found for males, but not for females (Table 2). Haplotype C was associated with decreased male total fitness while haplotype D was associated with increased total fitness (Figure 1B).

**Figure 1.**
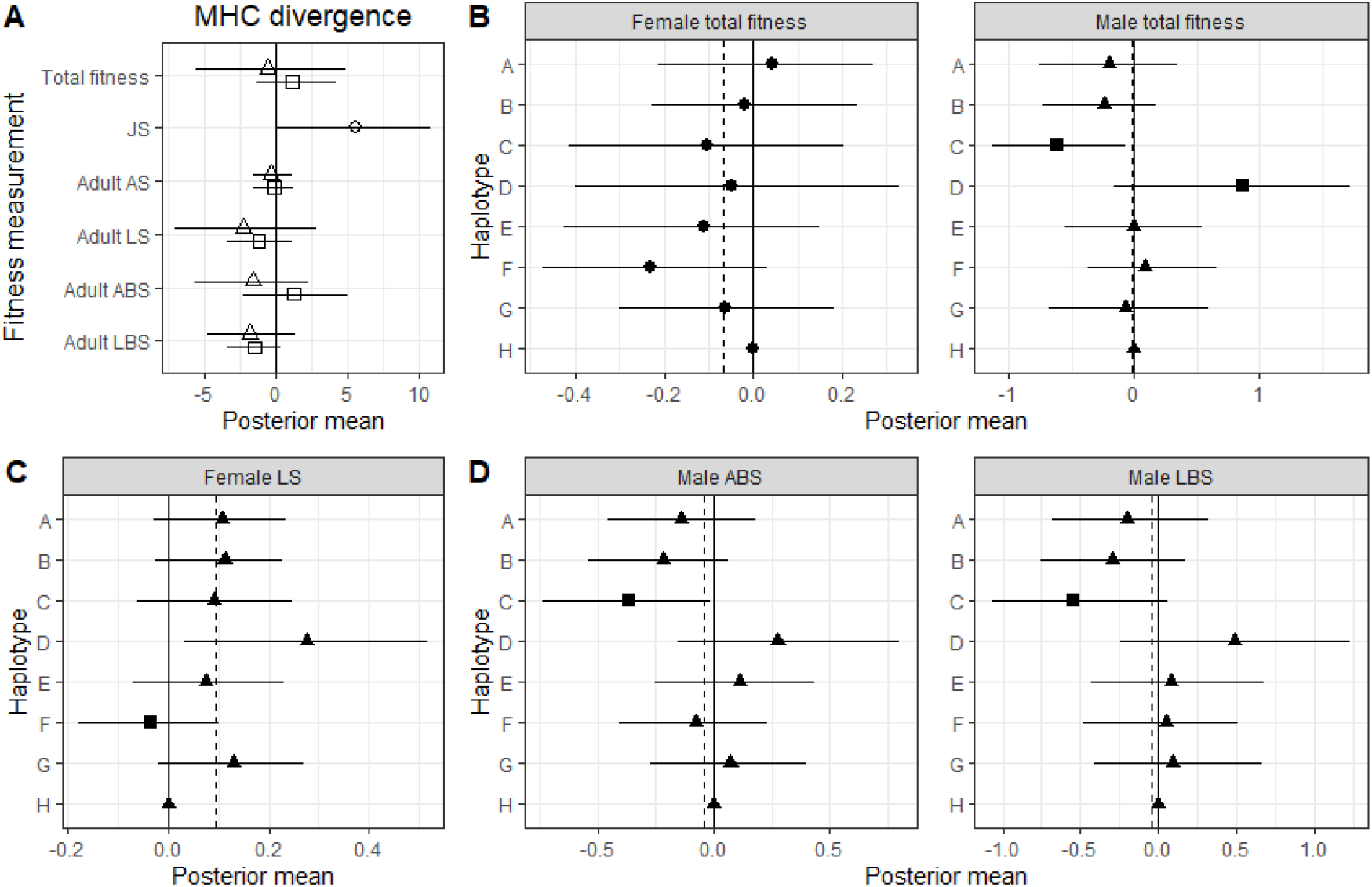
Associations between MHC and fitness measurements in Soay sheep derived from animal models. (A) Associations between MHC divergence and fitness measurements in juveniles (circle), females (squares) and males (triangles). Bars represent the 95% credibility intervals and those not overlapped with zero represent significant results. (B) Associations between MHC haplotypes and total fitness in females and males. The solid line represents the model intercept at haplotype H and posterior means and credible intervals for each haplotype are plotted relative to H. Dashed lines indicate the average posterior mean of all eight haplotypes. Circles indicate the Wald test was not significant, other symbols indicate it was significant. Haplotype effects that were significantly different from the mean are shown as squares with non-significant ones show as triangles. (C and D) Associations between MHC haplotypes and adult fitness components in females (C) and males (D). Symbols and lines as for B.

**Table 2.** Results (p value) of Wald tests for animal models testing for differences between haplotypes in fitness measures (d.f.=7). Bold numbers show significant results (p <0.05).

| Sex | Fitness measurement | p value |
| --- | --- | --- |
| / | JS | 0.13 |
| Female | Total fitness | 0.45 |
|  | AS | 0.073 |
|  | **LS** | **0.028** |
|  | ABS | 0.92 |
|  | LBS | 0.12 |
| Male | **Total fitness** | **0.0034** |
|  | AS | 0.39 |
|  | LS | 0.55 |
|  | **ABS** | **0.019** |
|  | **LBS** | **0.039** |

#### Juvenile survival

We found a positive association between MHC divergence and JS (Figure 1A). Considering haplotype effects, the Wald test was not significant (Table 2).

#### Adult annual survival

There was no association between MHC heterozygosity or MHC divergence and adult annual survival (AS) in either sex (Figure 1A). The Wald test for haplotype effects in AS was not significant for either sex (Table 2).

#### Adult life span

There was no association between MHC heterozygosity or MHC divergence and adult life span (LS) in either sex (Figure 1A). In the Wald test for haplotype effects a significant association was found for females, but not for males (Table 2). Haplotype F was associated with decreased adult female life span (Figure 1C).

#### Annual breeding success

There was no association between MHC heterozygosity or MHC divergence and annual breeding success (ABS) in either sex (Figure 1A). In the Wald test for haplotype effects a significant association was found for males, but not for females (Table 2). Haplotype C was associated with decreased adult male annual breeding success (Figure 1D).

#### Lifetime breeding success

We found no association between MHC heterozygosity or MHC divergence and adult lifetime breeding success (LBS) in either sex (Figure 1A). In the Wald test for haplotype effects a significant association was found for males, but not for females (Table 2). Haplotype C were associated with decreased adult male lifetime breeding success (Figure 1D).

#### Models without fitting the pedigree

The results obtained from models not including the pedigree were generally consistent with those above for associations between MHC heterozygosity/divergence and fitness measurements (Supplementary 5). For associations between individual MHC haplotypes and fitness measurements, estimates were mostly similar, but there were minor variations in terms of significance of individual haplotypes (Table 1, Supplementary 5). One difference was that we that found haplotype D was associated with increased female adult LS in models without pedigree fitted (Supplementary 5).

### Gene-drop analysis

All haplotype frequency changes were within expectation due to drift alone except for haplotype D (Figure 2). Of the 5000 simulations for Haplotype D, only 97 simulated regression slopes were greater than the slope generated from real data (*p*=0.0194). Therefore, this haplotype increased in frequency more than predicted by chance.

**Figure 2.**
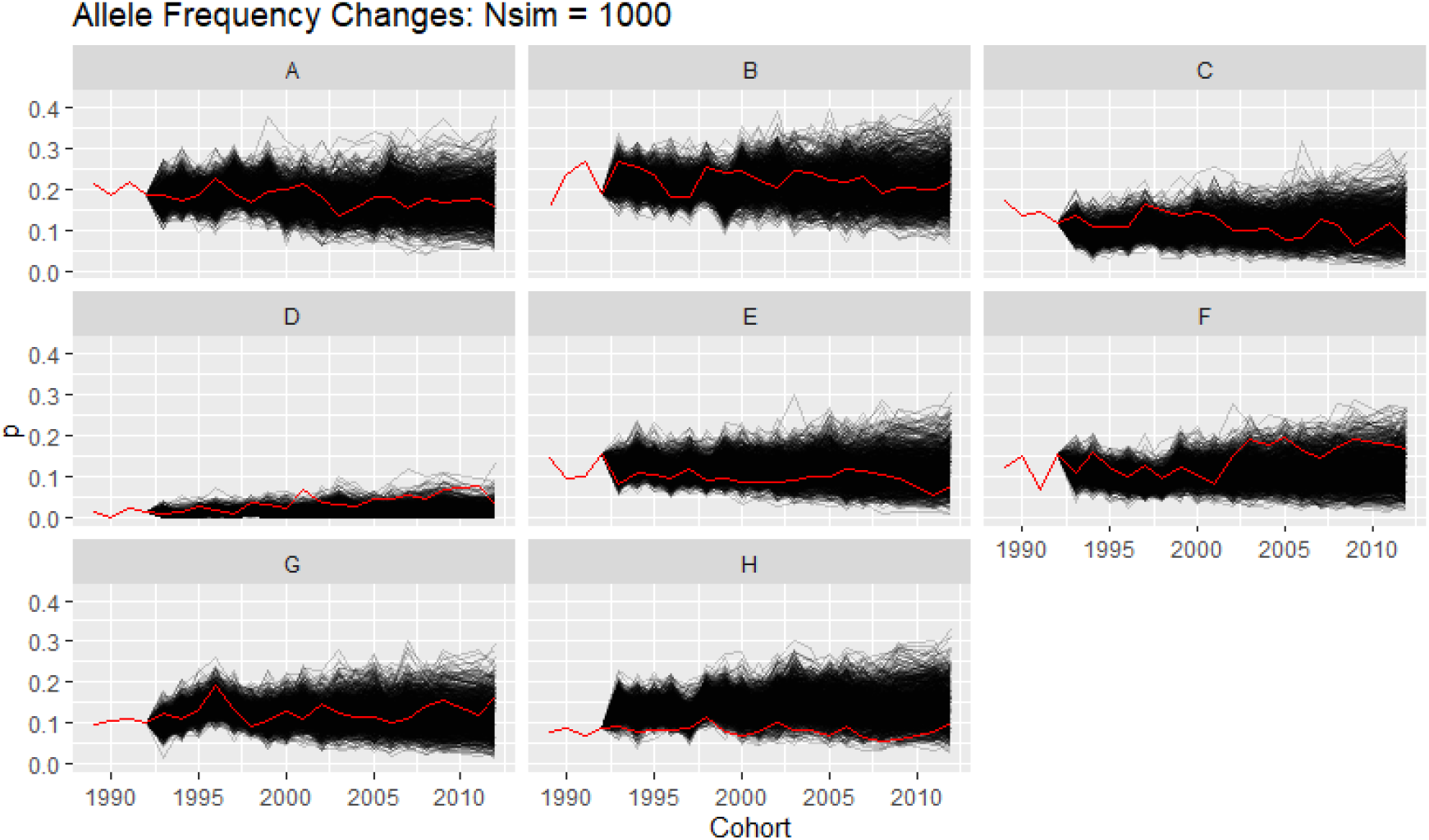
Results of the gene drop analysis for each haplotype. The black lines represent the result of simulations (N=5000) and the red lines represent the observed frequency of each haplotype in the Soay sheep population over the same time interval.

## Discussion

In this study we investigated associations between MHC class II diversity (heterozygosity, divergence and specific haplotypes) and fitness measurements including total fitness and five fitness components in a sample of individually-monitored free-living ruminants using generalized linear mixed models. We fitted MHC divergence, MHC heterozygosity and individual MHC haplotype effects together to test whether there is divergent allele/heterozygote advantage, rare-allele advantage or fluctuation selection. For all models, we included genetic relatedness to eliminate potential false positive results due to other genomic regions in common in related individuals and a genome inbreeding estimator to control genome-wide heterozygosity and the effect of inbreeding depression. We found a negative association between haplotype C and male total fitness and a positive association between haplotype D and male total fitness. We found MHC divergence was associated with increased JS. We also identified a negative association between haplotype C and adult male ABS and LBS and a negative association between haplotype F and adult female LS. Finally, we found that the rarest haplotype, D, increased significantly in frequency over the 23 year study period.

In relation to the various hypotheses for the evolution of MHC diversity, our study supports multiple hypotheses. In wild populations, evidence of MHC heterozygote fitness advantage has been reported in some previous studies (28, 29, 35) but not in others (24, 58). In addition, some studies have found evidence of divergent allele advantage, over and above heterozygote advantage, for example (40). In this study we found evidence for the divergent allele advantage hypothesis as MHC divergence was positively associated with JS independent of MHC heterozygosity. In addition, selection favouring or disfavouring particular haplotypes, as is likely under negative frequency-dependent selection when not in equilibrium or by the fluctuating selection hypothesis, has been reported in previous studies (31, 44). In this study, we also found evidence for negative frequency-dependent selection or fluctuating selection in that specific MHC haplotypes were associated with fitness measurements including total fitness, adult female LS and adult male ABS and LBS. Although haplotype D is rare, favoured and increasing, suggestive of rare allele advantage, our results are consistent with both rare allele advantage and fluctuating selection andwe are not capable of differentiating them at this time (13).

In this study, we found that MHC-fitness associations varied among different fitness measurements. We found evidence of divergent allele advantage in juvenile survival, but did not observe any evidence of these effects in total fitness and adult fitness components. These results are consistent with a previous study of Seychelles warbler that found a positive association between MHC diversity and juvenile but not adult survival (35). In Soay sheep, parasite infection intensity (measured as strongyle egg count) (59) and mortality is highest in the juvenile period (60) and parasite infection intensity is also negatively correlated with juvenile survival (61). Therefore, lambs with divergent MHC haplotypes may be better able to respond to a wider range of pathogen antigens, increasing their chances of surviving the harsh juvenile period. However, as more MHC homozygote lambs die in their first year, selection for divergent MHC constitution may no longer be evident in adults due to selective disappearance. Alternatively, the lack of selection for MHC heterozygosity or divergence in adult sheep could also result from maturation of the immune system or a change in the parasite community (59, 62). In terms of specific haplotypes, we did not find any association between specific MHC haplotypes and juvenile survival, but we did detect associations with total fitness and adult fitness components which could be a reflection of the heterogeneity of parasite infection and the development of immunity to parasites acquired between the juvenile and adult periods.

Although many studies in wild populations have identified associations between specific MHC alleles and fitness, few studies have observed an allele frequency change in response to selection. One exception is a study of the great reed warbler (Acrocephalus arundinaceus) that identified significantly higher variation in MHC allele frequencies between cohorts than at neutral loci, suggesting that fluctuating selection may be acting on MHC variation (63). In a previous experimental study, MHC alleles providing resistance to the respective specific parasite increased in frequency in the next host generation (64). If MHC-linked fitness effects are strong enough, we should be able to see changes in haplotype frequency over time in Soay sheep. In our study, we found haplotype C and D were associated with decreased and increased male total fitness respectively. In addition, the point estimates of haplotype C and D for female total fitness are, if anything, in the same direction (though not significant). Consistent with this, the gene drop analysis showed that haplotype D has increased significantly through the study period (p=0.0194). However, the frequency of haplotype C did not change more than expected by chance. A possible reason for this difference lies in the effects on adult life span (see Figure 1B, supplementary figure s5.2). Haplotype C was not significantly associated with adult life span in either sex, whereas haplotype D was associated with adult female life span and the point estimates of haplotype D for male adult life span was in the same direction though not significant. If individuals carrying haplotype D remain in the population longer, as this suggests, then they contribute to the significant increase in haplotype frequency in standing population each year. To our knowledge, our study is the first demonstrating that the frequency of a MHC haplotype conferring selective advantage also increased through the study period in a wild population.

Several aspects of our analysis improve on earlier MHC-fitness association studies. Benefiting from MHC haplotype data generated by locus-specific genotyping, this is one of the few studies using dosage models to precisely examine whether there is evidence for heterozygote advantage and divergent allele advantage which are caused by non-additive genetic effects (65, 66). Indeed, we demonstrated signature of divergent haplotype advantage in JS. In addition, to our knowledge, this is the first study to use animal models in a study of association between MHC and fitness measurements in wild populations. As the AM accounts for relatedness, we predicted that it would be more conservative and show fewer associations than GLMMs that did not account for relatedness. In fact, we found the results to be largely consistent between the two models. This may be due to the generally low heritability of the fitness traits (Supplementary 9). However, for traits which have a higher heritability, the results of animal models will be more conservative as they would exclude the false positive effects caused by genomic regions other than the MHC.

Long term individual-based studies like the Soay sheep project also enable repeat measurements of the strength and direction of selection over time (67, 68). In a previous study, three MHC-linked microsatellite alleles were found to be associated with survival in Soay sheep using individuals from 1985 to 1994 (44). Microsatellite alleles OLADRB 257 and 205 were associated with decreased juvenile and yearling survival respectively whereas the OLADRB 263 allele was associated with increased yearling survival. Some microsatellite alleles can be linked directly to the MHC class II haplotypes studied here: OLADRB allele 257 is linked to haplotype B (Supplementary 10). Thus, we might expect some consistency between the results from the previous study and the current one. However, in our analysis of juvenile survival, we did not find any evidence for haplotype-specific effects (Wald tests not significant; Table 2). The inconsistency between the previous and current study is probably due to differences in genotyping method, sample size and statistical methods including fitting inbreeding. In addition, alleles OLADRB 205 and 263 are associated with multiple haplotypes, and we have studied annual survival rather than yearling survival, which may also contribute to a failure to repeat the results.

Here we have identified a number of associations between MHC diversity and fitness measurements which are likely due to PMS. We would therefore expect to observe similar associations between MHC class II variation and measures of immunity to key parasites. In a previous study using the same MHC haplotype dataset, we examined associations between MHC class II variation and five phenotypic traits including strongyle faecal egg count, FEC and titres of Teladorsagia circumcincta specific immunoglobulin isotypes IgA, IgE and IgG. Although FEC is negatively correlated with fitness, no significant association between MHC variation and FEC was identified. In contrast, MHC divergence was found to be positively associated with lamb IgA titres. Specific haplotypes were also associated with antibody titres including a positive association between haplotype C and adult IgE and a positive association between haplotype D and IgE. However, no association was identifiedwith haplotype F that might match the female adult LS result found here. Our failure to identify MHC-FEC associations may result from the nature of this measure, its relationship with parasite burden and immunity to gastrointestinal nematode parasites. Other microorganisms, pathogenic or benign, are likely be involved in selection on MHC genes, including gut microbiota (69). Further studies examining associations between MHC diversity and the range of pathogens each animal is exposed to will help to clarify the nature of the selective forces acting to maintain MHC diversity in Soay sheep.

In summary, we used well-characterized MHC class II haplotypes to investigate MHC-linked fitness effects in Soay sheep. To our knowledge, our dataset, with more than 3000 individuals, is the largest yet used to study selection on MHC variation in a wild population. Our results support the existence of contemporary selection on MHC class II variation in Soay sheep, when genetic relatedness and inbreeding are controlled for. However, the selection mechanisms differs between juveniles and adults. We observed that the frequency of a rare MHC haplotype with a selective advantage has increased significantly during the study period. Our study highlights the importance of investigating selection on MHC genes across the whole lifespan and suggests different selection mechanisms on MHC variation could act simultaneously in a natural population.

## Acknowledgments

We thank the National Trust for Scotland for permission to work on St. Kilda and QinetiQ for logistics and support. We thank J. Pilkington, I. Stevenson and many volunteers who have helped with fieldwork on the island and all those who have contributed to keeping the project going over many years. W. Huang was supported by Edinburgh Global research scholarship. K. Ballingall received funding from the European Union’s Horizon 2020 research and innovation programme under grant agreement no. 731014KB (VetBioNet) and acknowledges the support received from the Scottish Government’s strategic research programme. The long-term project on St Kilda, the KASP genotyping and the SNP array genotyping were funded by the UK Natural Environment Research Council and the European Research Council.

## References

1. Obbard DJ, Welch JJ, Kim KW, & Jiggins FM (2009) Quantifying Adaptive Evolution in the Drosophila Immune System. Plos Genet 5(10).

2. Acevedo-Whitehouse K & Cunningham AA (2006) Is MHC enough for understanding wildlife immunogenetics? Trends Ecol Evol 21(8):433–438.

3. Bernatchez L & Landry C (2003) MHC studies in nonmodel vertebrates: what have we learned about natural selection in 15 years? Journal of Evolutionary Biology 16(3):363–377.

4. Piertney SB & Oliver MK (2006) The evolutionary ecology of the major histocompatibility complex. Heredity 96(1):7–21.

5. Garrigan D & Hedrick PW (2003) Detecting adaptive molecular polymorphism, lessons from the MHC. Am J Hum Genet 73(5):375–375.

6. Apanius V, Penn D, Slev PR, Ruff LR, & Potts WK (1997) The nature of selection on the major histocompatibility complex. Crit Rev Immunol 17(2):179–224.

7. Maccari G, et al. (2017) IPD-MHC 2.0: an improved inter-species database for the study of the major histocompatibility complex. Nucleic Acids Res 45(D1):D860–D864.

8. Oppelt C, Starkloff A, Rausch P, Von Holst D, & Rodel HG (2010) Major histocompatibility complex variation and age-specific endoparasite load in subadult European rabbits. Molecular ecology 19(19):4155–4167.

9. Radwan J, Kuduk K, Levy E, LeBas N, & Babik W (2014) Parasite load and MHC diversity in undisturbed and agriculturally modified habitats of the ornate dragon lizard. Molecular ecology 23(24):5966–5978.

10. Oliver MK, Telfer S, & Piertney SB (2009) Major histocompatibility complex (MHC) heterozygote superiority to natural multi-parasite infections in the water vole (Arvicola terrestris). Proceedings. Biological sciences 276(1659):1119–1128.

11. Westerdahl H, Asghar M, Hasselquist D, & Bensch S (2012) Quantitative disease resistance: to better understand parasite-mediated selection on major histocompatibility complex. Proceedings. Biological sciences 279(1728):577–584.

12. Jordan WC & Bruford MW (1998) New perspectives on mate choice and the MHC. Heredity 81:239–245.

13. Spurgin LG & Richardson DS (2010) How pathogens drive genetic diversity: MHC, mechanisms and misunderstandings. P R Soc B 277(1684):979–988.

14. Takahata N & Nei M (1990) Allelic Genealogy under Overdominant and Frequency-Dependent Selection and Polymorphism of Major Histocompatibility Complex Loci. Genetics 124(4):967–978.

15. Wakeland EK, et al. (1990) Ancestral polymorphisms of MHC class II genes: divergent allele advantage. Immunol Res 9(2):115–122.

16. Hughes AL & Nei M (1988) Pattern of Nucleotide Substitution at Major Histocompatibility Complex Class-I Loci Reveals Overdominant Selection. Nature 335(6186):167–170.

17. Ejsmond MJ & Radwan J (2015) Red Queen Processes Drive Positive Selection on Major Histocompatibility Complex (MHC) Genes. Plos Comput Biol 11(11).

18. Borghans JAM, Beltman JB, & De Boer RJ (2004) MHC polymorphism under host-pathogen coevolution. Immunogenetics 55(11):732–739.

19. Hedrick PW (2002) Pathogen resistance and genetic variation at MHC loci. Evolution; international journal of organic evolution 56(10):1902–1908.

20. Hill AVS, et al. (1991) Common West African Hla Antigens Are Associated with Protection from Severe Malaria. Nature 352(6336):595–600.

21. Rivero-de Aguilar J, et al. (2016) MHC-I provides both quantitative resistance and susceptibility to blood parasites in blue tits in the wild. J Avian Biol 47(5):669–677.

22. Westerdahl H, Stjernman M, Raberg L, Lannefors M, & Nilsson JA (2013) MHC-I Affects Infection Intensity but Not Infection Status with a Frequent Avian Malaria Parasite in Blue Tits. Plos One 8(8).

23. Westerdahl H, et al. (2005) Associations between malaria and MHC genes in a migratory songbird. P R Soc B 272(1571):1511–1518.

24. Radwan J, et al. (2012) MHC diversity, malaria and lifetime reproductive success in collared flycatchers. Molecular ecology 21(10):2469–2479.

25. Bolnick DI & Stutz WE (2017) Frequency dependence limits divergent evolution by favouring rare immigrants over residents. Nature 546(7657):285–+.

26. Phillips KP, et al. (2018) Immunogenetic novelty confers a selective advantage in host-pathogen coevolution. Proceedings of the National Academy of Sciences of the United States of America 115(7):1552–1557.

27. Sutton JT, Nakagawa S, Robertson BC, & Jamieson IG (2011) Disentangling the roles of natural selection and genetic drift in shaping variation at MHC immunity genes. Mol Ecol 20(21):4408–4420.

28. Dunn PO, Bollmer JL, Freeman-Gallant CR, & Whittingham LA (2013) Mhc Variation Is Related to a Sexually Selected Ornament, Survival, and Parasite Resistance in Common Yellowthroats. Evolution 67(3):679–687.

29. Banks SC, Dubach J, Viggers KL, & Lindenmayer DB (2010) Adult survival and microsatellite diversity in possums: effects of major histocompatibility complex-linked microsatellite diversity but not multilocus inbreeding estimators. Oecologia 162(2):359–370.

30. Lukasch B, et al. (2017) Major histocompatibility complex genes partly explain early survival in house sparrows. Sci Rep-Uk 7.

31. Sepil I, Lachish S, & Sheldon BC (2013) Mhc-linked survival and lifetime reproductive success in a wild population of great tits. Molecular ecology 22(2):384–396.

32. Kloch A, Baran K, Buczek M, Konarzewski M, & Radwan J (2013) MHC influences infection with parasites and winter survival in the root vole Microtus oeconomus. Evol Ecol 27(3):635–653.

33. Karlsson M, et al. (2015) House sparrowPasser domesticussurvival is not associated with MHC-I diversity, but possibly with specific MHC-I alleles. Journal of Avian Biology 46(2):167–174.

34. Pineaux M, et al. (2020) Sex and hatching order modulate the association between MHC-II diversity and fitness in early-life stages of a wild seabird. Mol Ecol 29(17):3316–3329.

35. Brouwer L, et al. (2010) MHC-dependent survival in a wild population: evidence for hidden genetic benefits gained through extra-pair fertilizations. Molecular ecology 19(16):3444–3455.

36. Babik W (2010) Methods for MHC genotyping in non-model vertebrates. Mol Ecol Resour 10(2):237–251.

37. Babik W, Taberlet P, Ejsmond MJ, & Radwan J (2009) New generation sequencers as a tool for genotyping of highly polymorphic multilocus MHC system. Mol Ecol Resour 9(3):713–719.

38. Osborne AJ, et al. (2015) Heterozygote advantage at MHC DRB may influence response to infectious disease epizootics. Molecular ecology 24(7):1419–1432.

39. Thoss M, Ilmonen P, Musolf K, & Penn DJ (2011) Major histocompatibility complex heterozygosity enhances reproductive success. Molecular ecology 20(7):1546–1557.

40. Lenz TL, Mueller B, Trillmich F, & Wolf JBW (2013) Divergent allele advantage at MHC-DRB through direct and maternal genotypic effects and its consequences for allele pool composition and mating. P R Soc B 280(1762).

41. Clutton-Brock T & Pemberton JM (2004) Soay sheep - Dynamics and selection in an island population. Cambridge University Press.

42. Berenos C, et al. (2015) Heterogeneity of genetic architecture of body size traits in a free-living population. Molecular ecology 24(8):1810–1830.

43. Paterson S (1998) Evidence for balancing selection at the major histocompatibility complex in a free-living ruminant. J Hered 89(4):289–294.

44. Paterson S, Wilson K, & Pemberton JM (1998) Major histocompatibility complex variation associated with juvenile survival and parasite resistance in a large unmanaged ungulate population (Ovis aries L.). Proceedings of the National Academy of Sciences of the United States of America 95(7):3714–3719.

45. Charbonnel N & Pemberton J (2005) A long-term genetic survey of an ungulate population reveals balancing selection acting on MHC through spatial and temporal fluctuations in selection. Heredity 95(5):377–388.

46. Dicks KL, Pemberton JM, & Ballingall KT (2019) Characterisation of major histocompatibility complex class IIa haplotypes in an island sheep population. Immunogenetics 71(5–6):383–393.

47. Dicks K (2017) Unravelling major histocompatibility complex diversity in the Soay Sheep of St.Kilda. University of Edinburgh.

48. Feulner PGD, et al. (2013) Introgression and the fate of domesticated genes in a wild mammal population. Molecular ecology 22(16):4210–4221.

49. Berenos C, Ellis PA, Pilkington JG, & Pemberton JM (2014) Estimating quantitative genetic parameters in wild populations: a comparison of pedigree and genomic approaches. Mol Ecol 23(14):3434–3451.

50. Huisman J (2017) Pedigree reconstruction from SNP data: parentage assignment, sibship clustering and beyond. Molecular Ecology Resources 17(5):1009–1024.

51. Morrissey MB, et al. (2012) The prediction of adaptive evolution: empirical application of the secondary theorem of selection and comparison to the breeder’s equation. Evolution; international journal of organic evolution 66(8):2399–2410.

52. Dicks KL, Pemberton JM, Ballingall KT, & Johnston SE (2020) Haplotyping MHC class IIa by high throughput screening in an isolated sheep population. bioRxiv:2020.2007.2020.212225.

53. Henikoff S (1996) Scores for sequence searches and alignments. Curr Opin Struc Biol 6(3):353–360.

54. Berenos C, Ellis PA, Pilkington JG, & Pemberton JM (2016) Genomic analysis reveals depression due to both individual and maternal inbreeding in a free-living mammal population. Mol Ecol 25(13):3152–3168.

55. Stoffel MA, Johnston SE, Pilkington JG, & Pemberton JM (2020) Genetic architecture and lifetime dynamics of inbreeding depression in a wild mammal. bioRxiv:2020.2005.2027.118877.

56. Hadfield JD (2010) MCMC Methods for Multi-Response Generalized Linear Mixed Models: The MCMCglmm R Package. J Stat Softw 33(2):1–22.

57. R Core Team (2013) R: A language and environment for statistical computing.

58. Sepil I, Lachish S, Hinks AE, & Sheldon BC (2013) Mhc supertypes confer both qualitative and quantitative resistance to avian malaria infections in a wild bird population. P R Soc B 280(1759).

59. Craig BH, Pilkington JG, & Pemberton JM (2006) Gastrointestinal nematode species burdens and host mortality in a feral sheep population. Parasitology 133:485–496.

60. Clutton-Brock TH & Pemberton JM (2004) Soay sheep: dynamics and selection in an island population (Cambridge University Press).

61. Hayward AD, et al. (2011) Natural selection on a measure of parasite resistance varies across ages and environmental conditions in a wild mammal. J Evolution Biol 24(8):1664–1676.

62. Sparks AM, et al. (2018) Natural Selection on Antihelminth Antibodies in a Wild Mammal Population. Am Nat 192(6):745–760.

63. Westerdahl H, Hansson B, Bensch S, & Hasselquist D (2004) Between-year variation of MHC allele frequencies in great reed warblers: selection or drift? J Evol Biol 17(3):485–492.

64. Eizaguirre C, Lenz TL, Kalbe M, & Milinski M (2012) Rapid and adaptive evolution of MHC genes under parasite selection in experimental vertebrate populations. Nature communications 3:621.

65. Hu XL, et al. (2015) Additive and interaction effects at three amino acid positions in HLA-DQ and HLA-DR molecules drive type 1 diabetes risk. Nature genetics 47(8):898–+.

66. Lenz TL, et al. (2015) Widespread non-additive and interaction effects within HLA loci modulate the risk of autoimmune diseases. Nature genetics 47(9):1085–+.

67. Siepielski AM, DiBattista JD, & Carlson SM (2009) It’s about time: the temporal dynamics of phenotypic selection in the wild. Ecol Lett 12(11):1261–1276.

68. Siepielski AM, DiBattista JD, Evans JA, & Carlson SM (2011) Differences in the temporal dynamics of phenotypic selection among fitness components in the wild. P R Soc B 278(1711):1572–1580.

69. Bolnick DI, et al. (2014) Major Histocompatibility Complex class IIb polymorphism influences gut microbiota composition and diversity. Molecular ecology 23(19):4831–4845.

