## Supplementary for "Age-dependent selection on MHC class 2 variation in a free-living ruminant"

**Supplementary 1 Frequency of MHC haplotypes in this study**

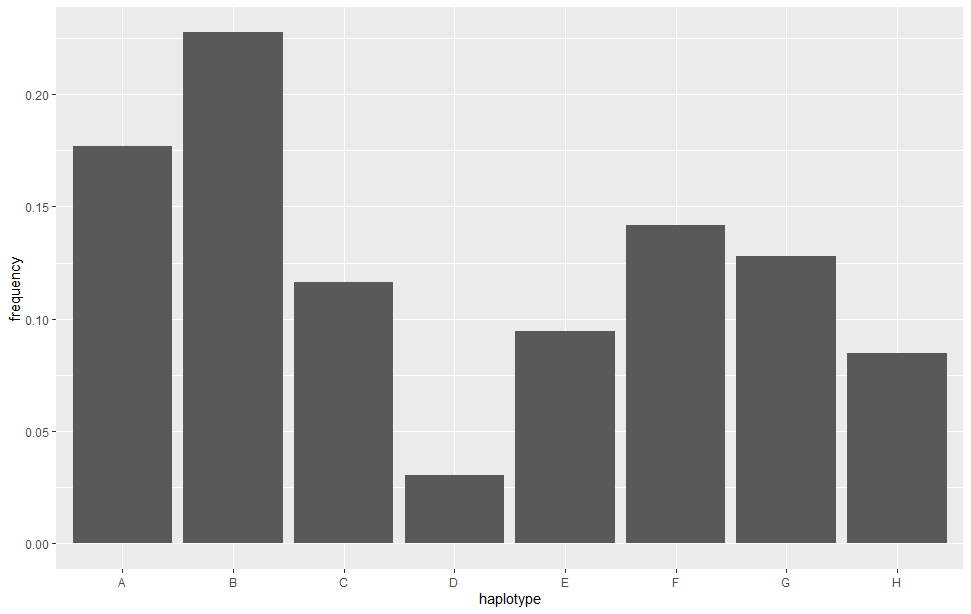

Figure S1.1. The frequency of each MHC haplotype in the sample of Soay sheep used in our study (N = 3440 individuals)

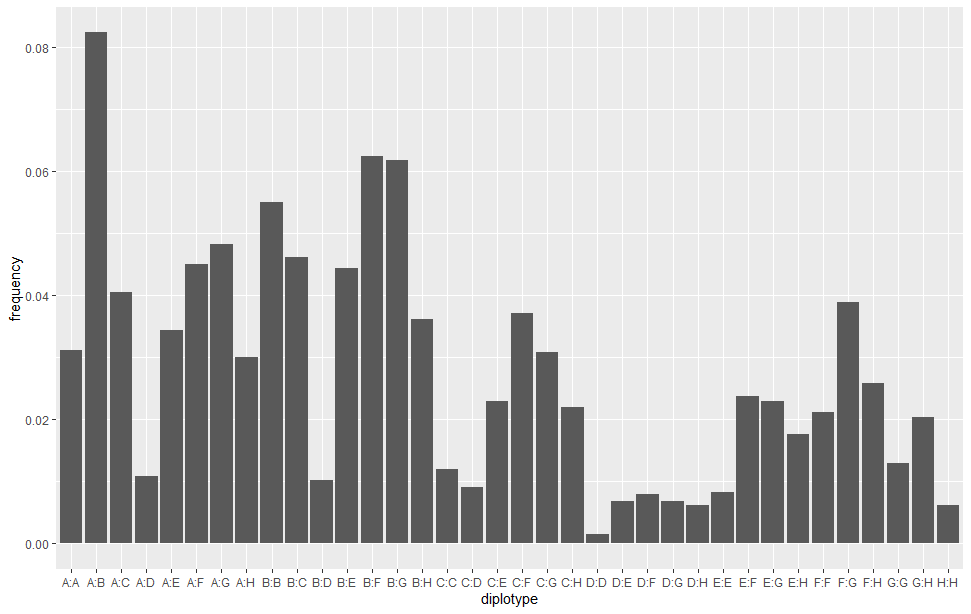

Figure S1.2. The frequency of each MHC diplotype in the sample of Soay sheep used in our study (N = 3440 individuals).

**Supplementary 2 MHC divergence**

Each MHC locus consists of a single allele except for haplotype G which has two alleles at DQB2. We used allele (DQB2*12*01) identified from a Soay RNA sequence to calculate p-distance. There are seven expressed loci identified in sheep but each haplotype only consists of five loci with either DRB1+DQA2+DQB2+DQA1+DQB1 or DRB1+DQA2+DQB2+DQA2-like+DQB2-like combinations. To tackle the problem of null alleles at certain loci, we aligned DQA1 alleles with DQA2-like alleles and DQB1 alleles with DQB2-like alleles. Thus, p-distance was calculated from an alignment composed of five loci for each haplotype.

Table S2. The proportion of the amino acid differences (p-distance) within exon 2 between MHC haplotypes.

|  | A | B | C | D | E | F | G | H |
| --- | --- | --- | --- | --- | --- | --- | --- | --- |
| A |  |  |  |  |  |  |  |  |
| B | 0.1462 |  |  |  |  |  |  |  |
| C | 0.2088 | 0.1624 |  |  |  |  |  |  |
| D | 0.2390 | 0.1671 | 0.1810 |  |  |  |  |  |
| E | 0.1907 | 0.1791 | 0.2605 | 0.2512 |  |  |  |  |
| F | 0.1949 | 0.1323 | 0.1926 | 0.1694 | 0.2279 |  |  |  |
| G | 0.1465 | 0.1651 | 0.2512 | 0.2651 | 0.0674 | 0.2093 |  |  |
| H | 0.1949 | 0.0974 | 0.1601 | 0.1879 | 0.2233 | 0.1392 | 0.2140 |  |

**Supplementary 3 Distribution of fitness components**

***Total fitness:***

There are large differences total fitness, measured as lifetime breeding success between females and males. The range in male breeding success is much larger than in females. As juveniles were included, there were a substantial of individuals with no offspring in both females and males. The maximum number of offspring in males was 108 while the maximum number of offspring in females was 20.

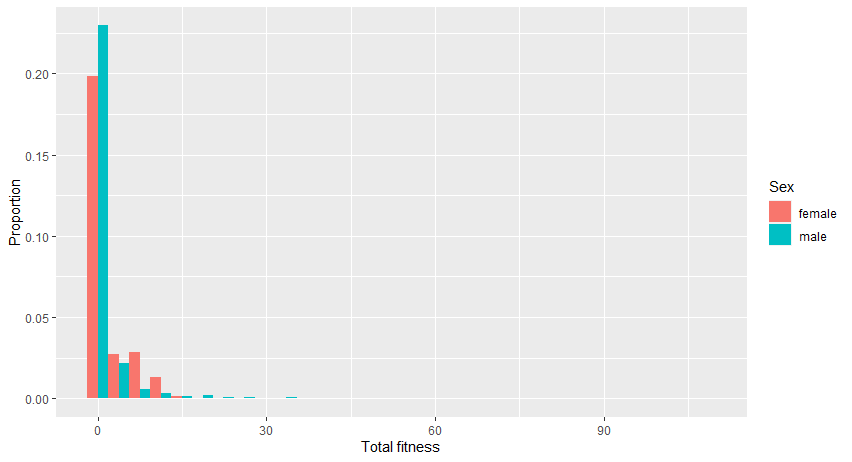

Figure S3.1.Distribution of total fitness of all the sheep used in our study (N=3440).

***Juvenile survival and life span:***

Most sheep (more than 60%) died during the juvenile period. Females had a slightly higher juvenile survival rate. After the juvenile period, the life span of Soay sheep also differed between females and males. Female sheep can live up to ~16 years of age, while male sheep can only live up to ~10 years, with most males dying before they are 5 years old.

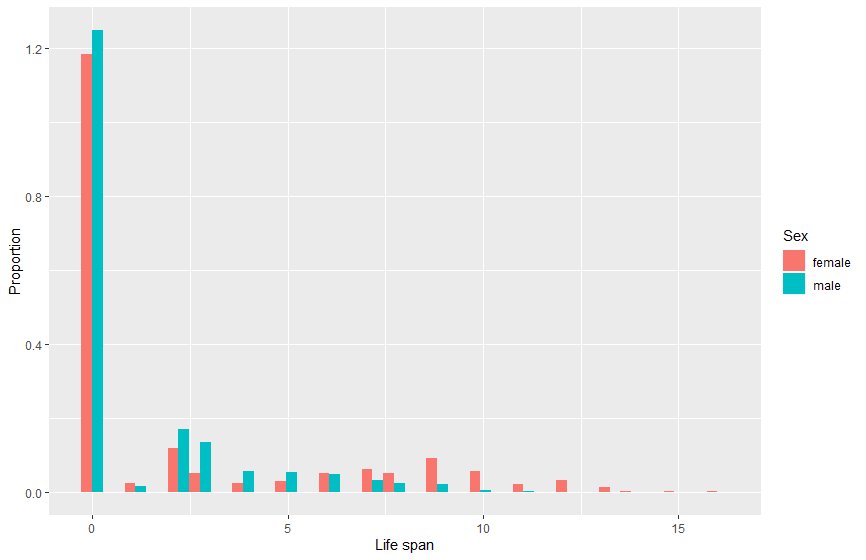

Figure S3.2. Distribution of life span of all the sheep used in our study (N=3440).

***Adult annual breeding success and lifetime breeding success:***

There are large differences in breeding success between females and males that survived at least to age one year. The range in male breeding success is much larger than in females. For ABS, males had up to 24 offspring per year, while females only had up to three offspring. For LBS, the maximum number of offspring in males was 108, with most males siring no offspring. Most females had one or more offspring, with a maximum number of 20 over their lifetime.

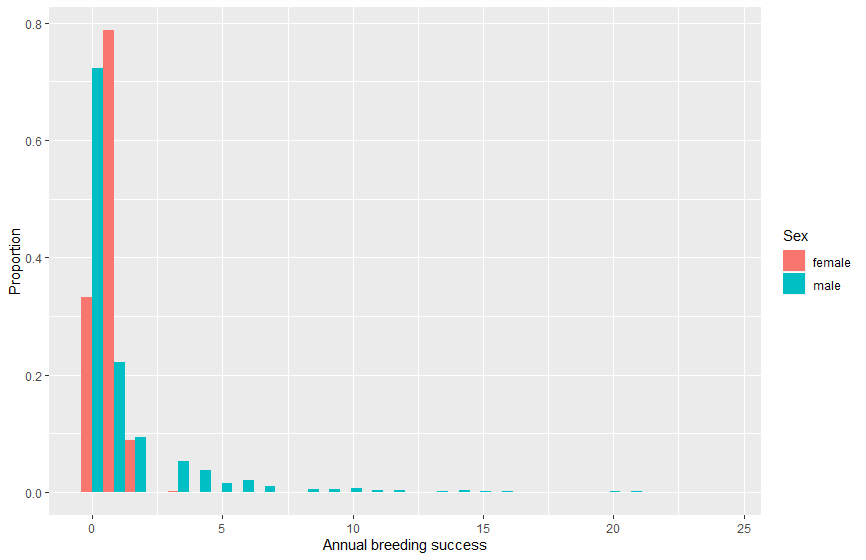

Figure S3.3 Distribution of adult annual breeding success of the sheep (N=1119).

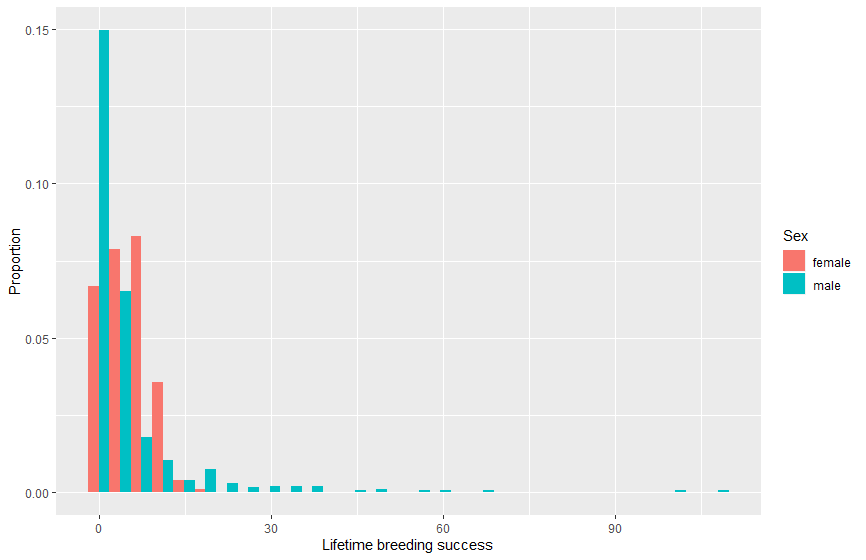

Figure S3.4 Distribution of adult lifetime breeding sucess of the sheep (N=1119).

**Supplementary 4 Additional tests to verify associations between fitness and individual haplotype**

The posterior mean and covariances of the haplotype effects were used to construct a Wald test to evaluate whether fitness measures differed systematically between individuals with different haplotypes. When Wald tests were ‘significant’ we conducted an additional test to confirm which haplotype(s) differed from the rest. We did this by obtaining the posterior distribution of the difference between the effect of a focal haplotype and the average effect of non-focal haplotypes. An MCMC p-value was calculated as 2p, where p is either the posterior probability that the difference is less than zero or greater than zero, whichever is smaller (1). The posterior probability is approximated by the proportion of MCMC samples that crossed zero.

**Supplementary 5 Associations between MHC haplotypes and fitness components in Soay sheep.**

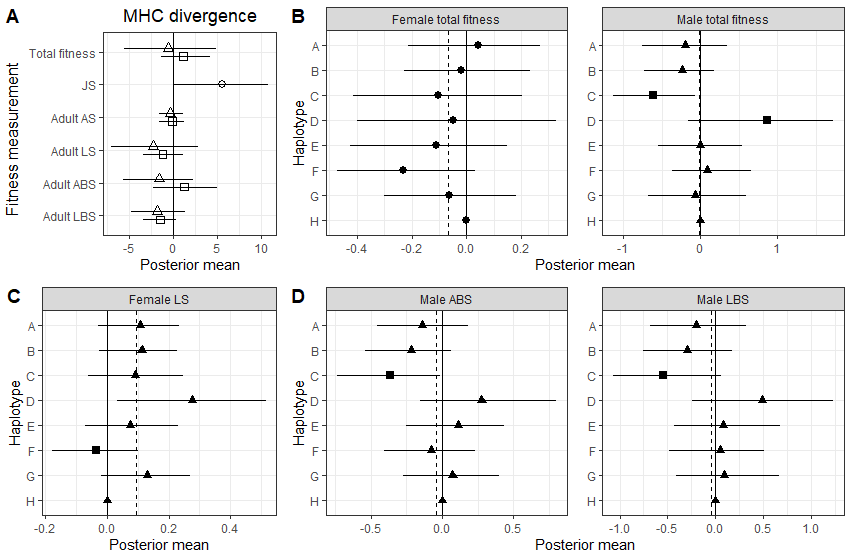

Figure S5.1. Associations between MHC and fitness measurements in Soay sheep derived from GLMMs that did not include the pedigree. (A) Associations between MHC divergence and fitness measurement in juveniles, (circles), adult females (squares) and adult males (triangles). Bars represent the 95% credibility intervals and those not overlapped with zero represent significant results. (B) Associations between MHC haplotypes and total fitness. The solid line represents the model intercept at haplotype H and posterior means and credible intervals for each haplotype are plotted relative to H. Dashed lines indicate the average posterior mean of all eight haplotypes. Circles suggests Wald test was not significant. Haplotype effects that were significantly different from the mean are shown as squares with non-significant ones presented in triangles if Wald test was significant. (C and D) Associations between MHC haplotypes and adult fitness components in females (C) and males (D). Symbols and lines as for B.

Table S5. Results (*p value*) of Wald tests for GLMMs (d.f.=7) . Bold numbers show significant results (*p* <0.05).

| Sex | Fitness component | p value |
| --- | --- | --- |
| / | JS | 0.096 |
| Female | Total fitness | 0.21 |
|  | AS | 0.064 |
|  | **LS** | **0.029** |
|  | ABS | 0.810 |
|  | LBS | 0.088 |
| Male | **Total fitness** | **0.029** |
|  | AS | 0.400 |
|  | LS | 0.640 |
|  | **ABS** | **0.014** |
|  | **LBS** | **0.046** |

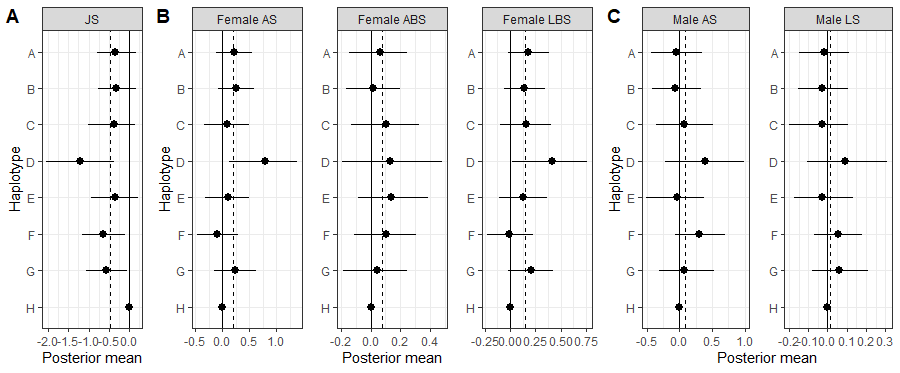

Figure S5.2. Associations between MHC haplotypes and fitness components in Soay sheep derived from animal models in juveniles (A), adult females (B) and adult males (C). Dashed lines indicate the average posterior mean of all eight haplotypes. Dashed lines indicate the average posterior mean of all eight haplotypes. Circles showed Wald tests of the models were not significant.

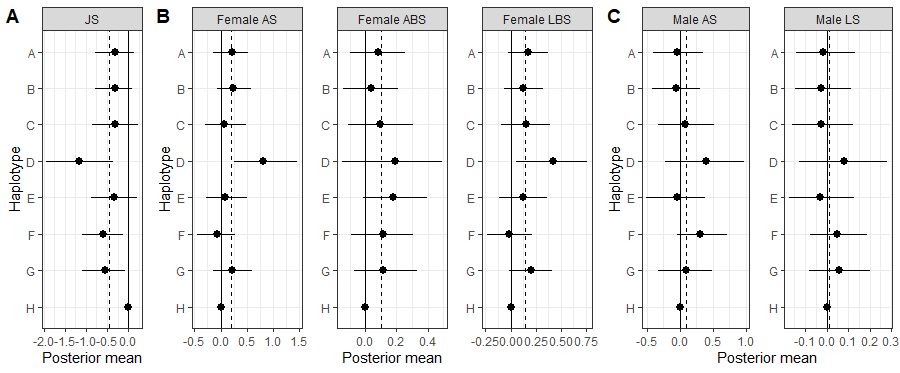

Figure S5.3. Associations between MHC haplotypes and fitness components in Soay sheep derived from GLMMs in juveniles (A), adult females (B) and adult males (C). Posterior means and credible intervals for each haplotypes were plotted in relative to haplotype H from original model outputs. Dashed lines indicate the average posterior mean of all eight haplotypes. Circles showed Wald tests of the models were not significant.

**Supplementary 6 Model outputs for fixed effects.**

Table S6.1 Results of total fitness model with pedigree fitted (Animal Models).

| Fixed effects | Male | | | Female | | |
| --- | --- | --- | --- | --- | --- | --- |
|  | Estimate | Lower 95 % CI | Upper 95 % CI | Estimate | Lower 95 % CI | Upper 95 % CI |
| Haplotype A | 0.042 | -0.213 | 0.270 | -0.194 | -0.755 | 0.352 |
| Haplotype B | -0.020 | -0.227 | 0.231 | -0.239 | -0.734 | 0.183 |
| Haplotype C | -0.104 | -0.412 | 0.205 | -0.612 | -1.126 | -0.061 |
| Haplotype D | -0.047 | -0.399 | 0.329 | 0.865 | -0.154 | 1.729 |
| Haplotype E | -0.110 | -0.425 | 0.147 | -0.005 | -0.550 | 0.546 |
| Haplotype F | -0.231 | -0.474 | 0.031 | 0.094 | -0.367 | 0.662 |
| Haplotype G | -0.064 | -0.300 | 0.183 | -0.072 | -0.684 | 0.595 |
| Fgrm | -6.426 | -9.613 | -2.917 | -13.167 | -18.299 | -7.003 |
| Heterozygosity | -0.174 | -0.655 | 0.362 | -0.481 | -1.404 | 0.354 |
| Divergence | 1.151 | -1.359 | 4.204 | -0.509 | -5.512 | 4.979 |

Table S6.2. Results of total fitness model without pedigree fitted (GLMMs).

| Fixed effects | Female | | | Male | | |
| --- | --- | --- | --- | --- | --- | --- |
|  | Estimate | Lower 95 % CI | Upper 95 % CI | Estimate | Lower 95 % CI | Upper 95 % CI |
| Haplotype A | 0.084 | -0.141 | 0.267 | -0.180 | -0.692 | 0.385 |
| Haplotype B | 0.011 | -0.189 | 0.217 | -0.241 | -0.730 | 0.307 |
| Haplotype C | -0.080 | -0.338 | 0.188 | -0.454 | -1.030 | 0.129 |
| Haplotype D | -0.008 | -0.374 | 0.369 | 0.678 | -0.127 | 1.512 |
| Haplotype E | -0.078 | -0.314 | 0.167 | 0.133 | -0.445 | 0.689 |
| Haplotype F | -0.187 | -0.436 | 0.043 | 0.229 | -0.282 | 0.773 |
| Haplotype G | 0.000 | -0.261 | 0.255 | -0.017 | -0.640 | 0.567 |
| Fgrm | -6.003 | -9.889 | -2.283 | -13.592 | -20.492 | -7.428 |
| Heterozygosity | -0.104 | -0.520 | 0.407 | -0.206 | -1.269 | 0.868 |
| Divergence | 0.680 | -1.779 | 2.901 | -1.563 | -6.763 | 3.876 |

Table S6.3. Results of juvenile survival model with pedigree fitted (Animal Models). Bold Font shows significant results effects of MHC divergence.

| Fixed effects | Estimate | Lower 95 % CI | Upper 95 % CI |
| --- | --- | --- | --- |
| Divergence | **5.657** | **0.069** | **10.873** |
| Heterozygosity | -0.576 | -1.649 | 0.369 |
| Haplotype A | -0.334 | -0.786 | 0.181 |
| Haplotype B | -0.319 | -0.771 | 0.198 |
| Haplotype C | -0.368 | -1.000 | 0.153 |
| Haplotype D | -1.218 | -2.058 | -0.372 |
| Haplotype E | -0.330 | -0.926 | 0.227 |
| Haplotype F | -0.632 | -1.161 | -0.099 |
| Haplotype G | -0.567 | -1.074 | -0.036 |
| Litter size | -0.543 | -0.693 | -0.405 |
| Fgrm | -3.250 | -5.031 | -1.186 |
| Sex | -0.437 | -1.025 | 0.146 |
| Divergence*Sex | -2.637 | -5.881 | 0.747 |
| Heterozygosity*Sex | 0.288 | -0.294 | 0.971 |
| Haplotype A*Sex | 0.180 | -0.124 | 0.501 |
| Haplotype B*Sex | 0.153 | -0.142 | 0.473 |
| Haplotype C*Sex | 0.118 | -0.227 | 0.474 |
| Haplotype D*Sex | 0.740 | 0.217 | 1.266 |
| Haplotype E*Sex | 0.119 | -0.279 | 0.459 |
| Haplotype F*Sex | 0.356 | 0.023 | 0.697 |
| Haplotype G*Sex | 0.247 | -0.092 | 0.568 |

Table S6.4. Results of juvenile survival model without pedigree fitted (GLMMs). Bold Font shows significant results effects of MHC divergence.

| Fixed effects | Estimate | Lower 95 % CI | Upper 95 % CI |
| --- | --- | --- | --- |
| Divergence | **5.052** | **0.068** | **10.150** |
| Heterozygosity | -0.479 | -1.412 | 0.500 |
| Haplotype A | -0.307 | -0.785 | 0.134 |
| Haplotype B | -0.323 | -0.781 | 0.087 |
| Haplotype C | -0.304 | -0.861 | 0.223 |
| Haplotype D | -1.156 | -1.938 | -0.365 |
| Haplotype E | -0.324 | -0.874 | 0.215 |
| Haplotype F | -0.605 | -1.089 | -0.126 |
| Haplotype G | -0.554 | -1.083 | -0.074 |
| Litter size | -0.549 | -0.688 | -0.421 |
| Fgrm | -2.994 | -4.802 | -1.047 |
| Sex | -0.414 | -0.958 | 0.129 |
| Divergence*Sex | -2.313 | -5.464 | 0.918 |
| Heterozygosity*Sex | 0.238 | -0.314 | 0.907 |
| Haplotype A*Sex | 0.165 | -0.137 | 0.460 |
| Haplotype B*Sex | 0.157 | -0.113 | 0.442 |
| Haplotype C*Sex | 0.105 | -0.226 | 0.447 |
| Haplotype D*Sex | 0.699 | 0.224 | 1.212 |
| Haplotype E*Sex | 0.119 | -0.229 | 0.462 |
| Haplotype F*Sex | 0.346 | 0.035 | 0.649 |
| Haplotype G*Sex | 0.231 | -0.081 | 0.569 |

Table S6.5. Results of models (adult fitness components) with pedigree fitted (Animal Models). Bold Font shows significant results in Wald tests (for haplotypes).

| Fixed effects | Sex | Female | | | | Male | | | |
| --- | --- | --- | --- | --- | --- | --- | --- | --- | --- |
|  | Fitness components | AS | **LS** | ABS | LBS | AS | LS | **ABS** | **LBS** |
| Age | Estimate | 0.132 |  | 0.569 |  | 0.380 |  | **0.961** |  |
|  | Lower 95 % CI | -0.013 |  | 0.506 |  | 0.055 |  | **0.846** |  |
|  | Upper 95% CI | 0.255 |  | 0.625 |  | 0.693 |  | **1.081** |  |
| Age2 | Estimate | -0.034 |  | -0.042 |  | -0.075 |  | **-0.061** |  |
|  | Lower 95 % CI | -0.043 |  | -0.047 |  | -0.105 |  | **-0.073** |  |
|  | Upper 95% CI | -0.025 |  | -0.037 |  | -0.045 |  | **-0.050** |  |
| Fgrm | Estimate | -5.828 | **-2.691** | -2.404 | -4.601 | -12.271 | -3.395 | **-4.675** | **-12.003** |
|  | Lower 95 % CI | -10.796 | **-4.518** | -4.868 | -7.741 | -18.285 | -5.267 | **-9.084** | **-18.353** |
|  | Upper 95% CI | -0.920 | **-0.660** | 0.378 | -1.639 | -6.345 | -1.574 | **-0.127** | **-4.969** |
| A | Estimate | 0.207 | **0.108** | 0.062 | 0.182 | -0.051 | -0.017 | **-0.138** | **-0.202** |
|  | Lower 95 % CI | -0.125 | **-0.030** | -0.142 | -0.017 | -0.435 | -0.150 | **-0.454** | **-0.677** |
|  | Upper 95% CI | 0.543 | **0.233** | 0.246 | 0.390 | 0.348 | 0.113 | **0.180** | **0.327** |
| B | Estimate | 0.253 | **0.113** | 0.017 | 0.135 | -0.063 | -0.027 | **-0.218** | **-0.298** |
|  | Lower 95 % CI | -0.082 | **-0.026** | -0.169 | -0.058 | -0.427 | -0.155 | **-0.539** | **-0.757** |
|  | Upper 95% CI | 0.583 | **0.227** | 0.199 | 0.342 | 0.327 | 0.104 | **0.064** | **0.178** |
| C | Estimate | 0.085 | **0.092** | 0.104 | 0.160 | 0.066 | -0.030 | **-0.363** | **-0.544** |
|  | Lower 95 % CI | -0.339 | **-0.060** | -0.135 | -0.093 | -0.366 | -0.198 | **-0.738** | **-1.064** |
|  | Upper 95% CI | 0.501 | **0.247** | 0.323 | 0.408 | 0.516 | 0.105 | **-0.017** | **0.059** |
| D | Estimate | 0.791 | **0.275** | 0.133 | 0.417 | 0.391 | 0.091 | **0.276** | **0.488** |
|  | Lower 95 % CI | 0.123 | **0.034** | -0.194 | 0.021 | -0.227 | -0.104 | **-0.152** | **-0.240** |
|  | Upper 95% CI | 1.374 | **0.514** | 0.481 | 0.762 | 0.995 | 0.311 | **0.801** | **1.236** |
| E | Estimate | 0.099 | **0.075** | 0.140 | 0.129 | -0.044 | -0.028 | **0.112** | **0.085** |
|  | Lower 95 % CI | -0.319 | **-0.070** | -0.086 | -0.109 | -0.504 | -0.173 | **-0.252** | **-0.427** |
|  | Upper 95% CI | 0.494 | **0.230** | 0.389 | 0.368 | 0.383 | 0.131 | **0.437** | **0.675** |
| F | Estimate | -0.091 | **-0.035** | 0.104 | -0.010 | 0.292 | 0.054 | **-0.079** | **0.051** |
|  | Lower 95 % CI | -0.475 | **-0.176** | -0.114 | -0.226 | -0.068 | -0.070 | **-0.406** | **-0.480** |
|  | Upper 95% CI | 0.283 | **0.101** | 0.309 | 0.226 | 0.695 | 0.181 | **0.234** | **0.513** |
| G | Estimate | 0.242 | **0.131** | 0.041 | 0.212 | 0.070 | 0.061 | **0.073** | **0.094** |
|  | Lower 95 % CI | -0.149 | **-0.020** | -0.185 | -0.023 | -0.320 | -0.080 | **-0.275** | **-0.411** |
|  | Upper 95% CI | 0.616 | **0.270** | 0.249 | 0.422 | 0.528 | 0.211 | **0.397** | **0.663** |
| Heterozygosity | Estimate | -0.388 | **-0.070** | 0.209 | 0.092 | -0.019 | -0.086 | **0.038** | **-0.220** |
|  | Lower 95 % CI | -1.075 | **-0.327** | -0.149 | -0.351 | -0.745 | -0.342 | **-0.558** | **-1.095** |
|  | Upper 95% CI | 0.335 | **0.204** | 0.570 | 0.510 | 0.739 | 0.170 | **0.594** | **0.757** |
| Divergence | Estimate | 1.314 | **-0.086** | -1.463 | -1.150 | -1.589 | -0.270 | **-1.803** | **-2.206** |
|  | Lower 95 % CI | -2.259 | **-1.544** | -3.335 | -3.369 | -5.682 | -1.611 | **-4.732** | **-7.035** |
|  | Upper 95% CI | 5.037 | **1.264** | 0.417 | 1.203 | 2.359 | 1.135 | **1.422** | **2.914** |

Table S6.6. Results of models (adult fitness components) without pedigree fitted (GLMMs). Bold Font shows significant results in Wald tests (for haplotypes).

| Fixed effects | Sex | Female | | | | Male | | | |
| --- | --- | --- | --- | --- | --- | --- | --- | --- | --- |
|  | Fitness components | AS | **LS** | ABS | LBS | AS | LS | **ABS** | **LBS** |
| Age | Estimate | 0.126 |  | 0.559 |  | 0.380 |  | **0.953** |  |
|  | Lower 95 % CI | -0.011 |  | 0.500 |  | 0.051 |  | **0.844** |  |
|  | Upper 95% CI | 0.254 |  | 0.618 |  | 0.689 |  | **1.073** |  |
| Age2 | Estimate | -0.033 |  | -0.042 |  | -0.074 |  | **-0.061** |  |
|  | Lower 95 % CI | -0.042 |  | -0.047 |  | -0.104 |  | **-0.072** |  |
|  | Upper 95% CI | -0.024 |  | -0.037 |  | -0.043 |  | **-0.050** |  |
| Fgrm | Estimate | -6.040 | **-2.713** | -2.246 | -4.666 | -12.220 | -3.447 | **-5.206** | **-12.377** |
|  | Lower 95 % CI | -11.146 | **-4.622** | -4.957 | -7.660 | -18.780 | -5.264 | **-9.481** | **-19.353** |
|  | Upper 95% CI | -1.122 | **-0.775** | 0.447 | -1.413 | -6.441 | -1.505 | **-0.378** | **-5.680** |
| A | Estimate | 0.213 | **0.102** | 0.083 | 0.174 | -0.039 | -0.015 | **-0.143** | **-0.181** |
|  | Lower 95 % CI | -0.139 | **-0.026** | -0.097 | -0.024 | -0.416 | -0.144 | **-0.473** | **-0.675** |
|  | Upper 95% CI | 0.521 | **0.227** | 0.256 | 0.369 | 0.348 | 0.132 | **0.147** | **0.333** |
| B | Estimate | 0.244 | **0.103** | 0.035 | 0.123 | -0.056 | -0.025 | **-0.217** | **-0.282** |
|  | Lower 95 % CI | -0.061 | **-0.027** | -0.141 | -0.067 | -0.422 | -0.146 | **-0.498** | **-0.754** |
|  | Upper 95% CI | 0.584 | **0.224** | 0.209 | 0.325 | 0.301 | 0.111 | **0.081** | **0.180** |
| C | Estimate | 0.065 | **0.080** | 0.093 | 0.149 | 0.084 | -0.028 | **-0.401** | **-0.519** |
|  | Lower 95 % CI | -0.290 | **-0.058** | -0.112 | -0.098 | -0.332 | -0.159 | **-0.752** | **-1.120** |
|  | Upper 95% CI | 0.487 | **0.241** | 0.308 | 0.393 | 0.512 | 0.123 | **-0.048** | **0.013** |
| D | Estimate | 0.803 | **0.285** | 0.194 | 0.426 | 0.389 | 0.080 | **0.213** | **0.438** |
|  | Lower 95 % CI | 0.254 | **0.069** | -0.149 | 0.051 | -0.232 | -0.130 | **-0.225** | **-0.254** |
|  | Upper 95% CI | 1.451 | **0.508** | 0.492 | 0.762 | 0.979 | 0.282 | **0.711** | **1.208** |
| E | Estimate | 0.085 | **0.065** | 0.176 | 0.119 | -0.046 | -0.030 | **0.097** | **0.082** |
|  | Lower 95 % CI | -0.279 | **-0.079** | -0.015 | -0.119 | -0.508 | -0.173 | **-0.229** | **-0.445** |
|  | Upper 95% CI | 0.499 | **0.204** | 0.396 | 0.364 | 0.379 | 0.128 | **0.436** | **0.658** |
| F | Estimate | -0.071 | **-0.036** | 0.112 | -0.012 | 0.298 | 0.050 | **-0.059** | **0.072** |
|  | Lower 95 % CI | -0.456 | **-0.170** | -0.088 | -0.234 | -0.046 | -0.079 | **-0.340** | **-0.458** |
|  | Upper 95% CI | 0.278 | **0.100** | 0.306 | 0.208 | 0.708 | 0.187 | **0.255** | **0.490** |
| G | Estimate | 0.223 | **0.112** | 0.115 | 0.203 | 0.086 | 0.058 | **0.062** | **0.114** |
|  | Lower 95 % CI | -0.142 | **-0.024** | -0.069 | -0.020 | -0.326 | -0.082 | **-0.292** | **-0.438** |
|  | Upper 95% CI | 0.591 | **0.243** | 0.330 | 0.414 | 0.492 | 0.202 | **0.380** | **0.632** |
| Heterozygosity | Estimate | -0.346 | **-0.053** | 0.242 | 0.116 | 0.000 | -0.100 | **-0.040** | **-0.244** |
|  | Lower 95 % CI | -1.035 | **-0.321** | -0.134 | -0.282 | -0.706 | -0.350 | **-0.605** | **-1.144** |
|  | Upper 95% CI | 0.375 | **0.220** | 0.605 | 0.559 | 0.779 | 0.141 | **0.545** | **0.742** |
| Divergence | Estimate | 1.182 | **-0.143** | -1.595 | -1.241 | -1.607 | -0.200 | **-1.310** | **-1.933** |
|  | Lower 95 % CI | -2.328 | **-1.563** | -3.514 | -3.471 | -5.466 | -1.590 | **-4.256** | **-6.566** |
|  | Upper 95% CI | 4.996 | **1.270** | 0.490 | 1.020 | 2.086 | 1.076 | **1.771** | **3.044** |

**Supplementary 7 Results of additional tests that verify the significance of individual haplotypes. Only models where Wald tests were significant are shown; the mean of estimates of all eight haplotypes for each model are also shown. Significant p values are presented in bold font.**

Table S7.1. Results of additional tests that verify significance of individual haplotype in animal models.

| Sex | Fitness measurements | Haplotype | Estimates | Mean | *p* |
| --- | --- | --- | --- | --- | --- |
| Female | Adult LS | A | 0.108 | 0.095 | 0.741 |
|  |  | B | 0.113 |  | 0.697 |
|  |  | C | 0.092 |  | 0.955 |
|  |  | D | 0.275 |  | 0.053 |
|  |  | E | 0.075 |  | 0.670 |
|  |  | **F** | **-0.035** |  | **0.001** |
|  |  | G | 0.131 |  | 0.428 |
|  |  | H | 0.000 |  | 0.068 |
| Male | Adult ABS | A | -0.138 | -0.042 | 0.294 |
|  |  | B | -0.218 |  | 0.057 |
|  |  | **C** | **-0.363** |  | **0.008** |
|  |  | D | 0.276 |  | 0.088 |
|  |  | E | 0.112 |  | 0.174 |
|  |  | F | -0.079 |  | 0.696 |
|  |  | G | 0.073 |  | 0.298 |
|  |  | H | 0.000 |  | 0.719 |
|  | Adult LBS | A | -0.202 | -0.041 | 0.254 |
|  |  | B | -0.298 |  | 0.095 |
|  |  | **C** | **-0.544** |  | **0.005** |
|  |  | D | 0.488 |  | 0.062 |
|  |  | E | 0.085 |  | 0.501 |
|  |  | F | 0.051 |  | 0.525 |
|  |  | G | 0.094 |  | 0.415 |
|  |  | H | 0.000 |  | 0.798 |
|  | Total fitness | A | -0.194 | -0.020 | 0.290 |
|  |  | B | -0.239 |  | 0.133 |
|  |  | **C** | **-0.612** |  | **0.006** |
|  |  | **D** | **0.865** |  | **0.033** |
|  |  | E | -0.005 |  | 0.995 |
|  |  | F | 0.094 |  | 0.445 |
|  |  | G | -0.072 |  | 0.790 |
|  |  | H | 0.000 |  | 0.908 |

Table S7.2. Results of additional tests that verify significance of individual haplotypes in GLMMs.

| Sex | Fitness measurements | Haplotype | Estimates | Mean | *p* |
| --- | --- | --- | --- | --- | --- |
| Female | Adult LS | A | 0.102 | 0.089 | 0.744 |
|  |  | B | 0.103 |  | 0.725 |
|  |  | C | 0.080 |  | 0.851 |
|  |  | **D** | **0.285** |  | **0.018** |
|  |  | E | 0.065 |  | 0.592 |
|  |  | **F** | **-0.036** |  | **0.003** |
|  |  | G | 0.112 |  | 0.580 |
|  |  | H | 0.000 |  | 0.084 |
| Male | Adult ABS | A | -0.143 | -0.056 | 0.353 |
|  |  | B | -0.217 |  | 0.068 |
|  |  | **C** | **-0.401** |  | **0.001** |
|  |  | D | 0.213 |  | 0.132 |
|  |  | E | 0.097 |  | 0.172 |
|  |  | F | -0.059 |  | 0.964 |
|  |  | G | 0.062 |  | 0.296 |
|  |  | H | 0.000 |  | 0.628 |
|  | Adult LBS | A | -0.181 | -0.034 | 0.292 |
|  |  | B | -0.282 |  | 0.088 |
|  |  | **C** | **-0.519** |  | **0.007** |
|  |  | D | 0.438 |  | 0.084 |
|  |  | E | 0.082 |  | 0.521 |
|  |  | F | 0.072 |  | 0.451 |
|  |  | G | 0.114 |  | 0.412 |
|  |  | H | 0.000 |  | 0.851 |
|  | Total fitness | A | -0.180 | 0.019 | 0.180 |
|  |  | B | -0.241 |  | 0.084 |
|  |  | **C** | **-0.454** |  | **0.004** |
|  |  | **D** | **0.678** |  | **0.033** |
|  |  | E | 0.133 |  | 0.522 |
|  |  | F | 0.229 |  | 0.192 |
|  |  | G | -0.017 |  | 0.871 |

**Supplementary 8 Results of gene drop analysis**

Table S8. Results of gene drop analysis. For each haplotype in the 5,000 simulations we counted the number of regression slopes of frequency change that were lower or higher than the observed data.

| Haplotype | Slopes lower than observed | Slopes higher than observed |
| --- | --- | --- |
| A | 2715 | 2285 |
| B | 2414 | 2586 |
| C | 1654 | 3346 |
| D | 4903 | 97 |
| E | 1754 | 3246 |
| F | 4734 | 266 |
| G | 2882 | 2118 |
| H | 217 | 4783 |

**Supplementary 9 Original model outputs of random effects and heritability of fitness traits.**

Table S9. Estimates of variance components and heritability (on the latent scale) of each fitness components (h^2^) in animal models. The variance components differed between fitness components. We show birth year effect (VBY), additive genetic effect (VA), permanent environment effect (VPE), measurement year effect (VCY) and the residual variance (VR).

| Sex | Fitness measurement | VBY | VA | VPE | VCY | VR | h^2^ |
| --- | --- | --- | --- | --- | --- | --- | --- |
| / | JS | 1.10 (0.52,1.91) | 0.12 (0.02,0.24) |  |  | 1.00 (1.00,1.00) | 0.06 (0.01,0.11) |
| Female | AS | 0.30 (0.03,0.66) | 0.25 (0.00,0.60) | 0.80 (0.29,1.35) | 1.17 (0.49,2.11) | 1.00 (1.00,1.00) | 0.07 (0.00,0.17) |
|  | LS | 0.07 (0.02,0.14) | 0.06 (0.00,0.12) |  |  | 0.06 (0.00,0.12) | 0.30 (0.00,0.62) |
|  | ABS | 0.03 (0.00,0.07) | 0.20 (0.06,0.35) | 0.08 (0.00,0.19) | 0.17 (0.07,0.28) | 1.00 (1.00,1.00) | 0.14 (0.05,0.23) |
|  | LBS | 0.14 (0.05,0.26) | 0.09 (0.00,0.22) |  |  | 0.38 (0.24,0.52) | 0.15 (0.00,0.35) |
| Male | AS | 0.18 (0.00,0.47) | 0.08 (0.00,0.28) | 0.80 (0.17,1.47) | 2.65 (1.16,4.55) | 1.00 (1.00,1.00) | 0.02 (0.00,0.06) |
|  | LS | 0.05 (0.01,0.10) | 0.01 (0.00,0.03) |  |  | 0.01 (0.00,0.03) | 0.12 (0.00,0.38) |
|  | ABS | 0.10 (0.00,0.24) | 0.30 (0.04,0.57) | 0.38 (0.10,0.64) | 0.14 (0.05,0.26) | 0.30 (0.21,0.39) | 0.25 (0.04,0.46) |
|  | LBS | 0.41 (0.08,0.84) | 0.36 (0.00,0.87) |  |  | 2.36 (1.72,3.07) | 0.11 (0.00,0.27) |

**Supplementary 10 OLADRB alleles and MHC class II haplotypes in Soay sheep**

Table S10. Associations between OLADRB alleles as used by Paterson et al (2) and class II haplotypes as used in the present study. OLADRB alleles denoted ^*^ were previously found by Paterson et al (2) to be associated with decreased juvenile and yearling survival, and those denoted ^‡^ with yearling survival.

| OLADRB microsatellite allele | MHC class II haplotype |
| --- | --- |
| 205^*^ | F and H |
| 213 | E |
| 257^*^ | B |
| 263^‡^ | A and G |
| 267 | A and G |
| 276 | C |
| 287 | D |

SI References

1.	Baayen RH, Davidson DJ, & Bates DM (2008) Mixed-effects modeling with crossed random effects for subjects and items. *J Mem Lang* 59(4):390-412.

2.	Paterson S, Wilson K, & Pemberton JM (1998) Major histocompatibility complex variation associated with juvenile survival and parasite resistance in a large unmanaged ungulate population (Ovis aries L.). *Proceedings of the National Academy of Sciences of the United States of America* 95(7):3714-3719.
